# MicroFPGA: an affordable FPGA platform for microscope control

**DOI:** 10.1101/2022.06.07.495178

**Authors:** Joran Deschamps, Christian Kieser, Philipp Hoess, Takahiro Deguchi, Jonas Ries

## Abstract

Modern microscopy relies increasingly on microscope automation to improve throughput, ensure reproducibility or observe rare events. Automation requires in particular computer control of the important elements of the microscope. Furthermore, optical elements that are usually fixed or manually movable can be placed on electronically-controllable elements. In most cases, a central electronics board is necessary to generate the control signals they require and to communicate with the computer. Fur such tasks, Arduino microcontrollers are widely used due to their low cost and programming entry barrier. However, they are limiting in their performance for applications that require high-speed or multiple parallel processes. Field programmable gate arrays (FPGA) are the perfect technology for high-speed microscope control, as they are capable of processing signals in parallel and with high temporal precision. While plummeting price made the technology available to consumers, a major hurdle remains the complex languages used to configure them. In this work, we used an affordable FPGA, delivered with an open-source and friendly-to-use programming language, to create a versatile microscope control platform called MicroFPGA. It is capable of synchronously triggering cameras and multiple lasers following complex patterns, as well as generating various signals used to control microscope elements such as filter wheels, servomotor stages, flip-mirrors, laser power or acoustooptic modulators. MicroFPGA is open-source and we provide online Micro-Manager, Java, Python and LabVIEW libraries, together with blueprints and tutorials.

## Introduction

Microscope automation is a promising avenue for biological studies. Not only does it save time for researchers, but it also substantially increases the throughput of the microscopes, improving statistical power or enabling the observation of rare events (1). In order to automate microscopes, all elements usually moved by hand must become computer-controlled. This often equates to mounting optics on motorized stages or servomotors, adding electronically-controlled flip-mirrors and shutters, and using electronics to synchronize devices such as cameras and lasers. Furthermore, automation enables the implementation of new modalities on the microscope, thus increasing the flexibility of the imaging. For instance, in fluorescence microscopy, lasers are often triggered using a camera signal in order to avoid unnecessary bleaching of the sample outside of the camera acquisition time frame. Adding a layer of signal processing in between the camera and the lasers enables more complex triggering patterns, such as fast pulsing or interleaved illumination.

Often, a central electronic board communicates with the computer and delivers the control signals to the various elements in order to move them into position. Typical signals include transistor-transistor logic (TTL), pulse-width modulation (PWM) or analog signals. Furthermore, reading out analog signals is a useful addition to monitor the microscope state, such as its temperature or laser power. Because instruments are subject to change, reprogrammable electronics are almost always preferred to application specific integrated circuits, as the cost of a new circuit production and long design-testing cycles are not suitable for research projects.

In that respect, Arduino microcontrollers have the advantage of being affordable, easy to program, open-source and benefit from a large community. They are therefore at the heart of many open-source microscopy projects (2–7). An intrinsic limitation of microcontrollers is that they are most often based on a single CPU, and therefore cannot perform multiple tasks in parallel. Processes, such as generating signals to drive servomotors, compete in CPU resources, limiting time resolution in signal processing. Careful optimization and use of interrupts can help perform complex tasks without perceptible delays. However, this approach does not scale well with an increasing number of processes and high-speed signal generation. Nonetheless, Arduino boards are sufficient for a very wide range of applications, in particular when fast processing is not crucial.

When the tasks at hand are too complex for microcontrollers, another solution is field-programmable gate arrays (FPGA). FP-GAs are integrated circuits consisting of millions of logic blocks that can be configured to perform the required tasks. Because independent tasks are broken down into different logic block ensembles, signals are processed concurrently. The number of processes that can run independently is then only limited by their complexity and the size of the FPGA. Thus, FPGAs are preferable to microcontrollers for applications requiring fast processing and precise timing. They are, however, considerably more complex to program. They typically require hardware description language (HDL) and advanced optimization tools, such as those provided by their manufacturers, limiting their use to specialists. Nonetheless, FPGAs have become more accessible, with more online resources available and plummeting prices (8). National Instruments (NI) offers a LabVIEW compatible FPGA module, which has found important applications in advanced optical microscopy (9–14), synchronizing complex microscopes for high-speed imaging or processing. In particular, using LabVIEW simplifies the coding task thanks to its intuitive visual programming. The use of LabVIEW-programmed FPGAs is not limited to NI’s proprietary software, but can be integrated with C or Python applications using the provided interfaces. Recently, the NI FPGA has even been used within Micro-Manager (15), an open-source microscope control software (16). Yet, the FPGA module and software are aimed at high-end applications, with an unreasonable cost for simple electronics control and signal processing tasks.

Here, we present MicroFPGA, a low-cost platform for electronic control of microscope devices aimed at providing a toolbox for automating microscopes. MicroFPGA is based on an affordable FPGA (Au+ or Cu, Alchitry), which rely solely on open-source or freely available software. Alchitry’s FPGAs are delivered with their own development environment (AlchitryLabs, Alchitry) and can be configured with a comparatively friendly HDL language: Lucid (17). MicroFPGA includes a powerful camera-laser synchronization module and laser triggering system, and can generate widely used electronic signals, such as TTL, PWM or servo control signals. In addition, the Au+ FPGA boards can read multiple analog signals. We also provide blueprints for complementary electronic boards, allowing voltage conversion and the generation of analog output signals. MicroFPGA is compatible with Micro-Manager and the online repositories contain Java, Python and LabVIEW libraries, as well as a guide and tutorials on how to use and modify the code (18, 19).

## Results

MicroFPGA aims at expanding the flexibility of a microscope by enabling computer control of existing or new microscope elements. As such, it serves as intermediary between the computer and the microscope in order to control these elements or measure signals from the microscope (see Fig. 1a). MicroFPGA can generate several types of signals commonly used to control devices in microscopy (see Fig. 1b and Table 1). The platform consists of the source code required to configure the FPGA boards, two complementary custom electronic boards for analog output and voltage conversion, as well as libraries to communicate with the FPGA from Micro-Manager, Java, Python and LabVIEW. In the next sections, we detail the FPGA boards used in this work, the additional electronics and the various signals available to control devices: laser triggering, PWM/analog out, servo motors, TTL and analog read-out.

**Fig. 1.**
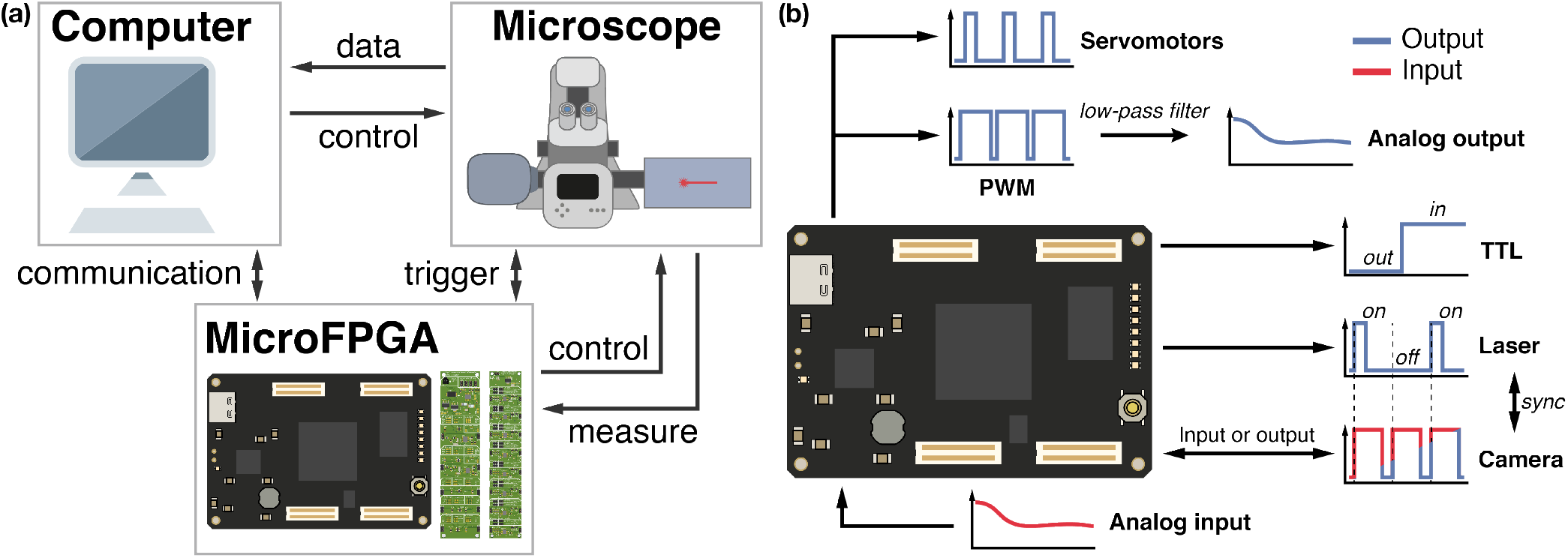
MicroFPGA overview. (a) MicroFPGA communicates with the computer and allows controlling or triggering elements on a microscope, as well as measuring signals. (b): Overview of the input and output signals of MicroFPGA.

**Table 1.** MicroFPGA inputs/outputs.

| Signal | I/O | Parameters | Board | # channels |
| --- | --- | --- | --- | --- |
| Camera | input | - | Au+/Cu | 1 |
| Camera | output | mode, start, pulse, delay, exposure, read-out | Au+/Cu | 1 |
| Laser | output | mode, duration, sequence | Au+/Cu | 8 |
| Analog | input | state | Au+ | 8 (fixed) |
| PWM | output | state | Au+/Cu | 5 |
| Servo | output | state | Au+/Cu | 7 |
| TTL | output | state | Au+/Cu | 4 |

### Alchitry FPGAs

Alchitry offers two different FPGA boards with 100 MHz on-board clocks (Au+ and Cu) at prices ranging from $50 (Cu) to $300 (Au+). The Au+ board is similar to the previous Au model and features an Artix 7 FPGA (Xilinx), which contains 101440 logic cells, a more powerful FPGA. The board also has an 12-bit analog-to-digital converter (ADC) capable of measuring sequentially analog voltages (0-1 V range) in 9 channels. In addition, it has 102 digital input/output (I/O) pins running at 3.3 V. Some pins can be configured to run at 1.8 V, but MicroFPGA uses only the higher voltage configuration. The Cu board is less powerful, with only 7680 logic elements in its iCE40-HX8K FPGA (Lattice) and 79 pins with 3.3 V I/O logic. Note that MicroFPGA is compatible with the previous Mojo and Au boards.

The source code for MicroFPGA has been written in Lucid (Alchitry) (17), a friendly HDL language developed specifically for Alchitry’s FPGA boards. A third-party tool is necessary in order to compile the FPGA code using AlchitryLabs (Alchitry). All Alchitry FPGAs can be compiled using proprietary tools available with free licenses: iCEcube2 (Lattice semiconductors) and Vivado (Xilinx) for the Cu and Au+, respectively. Alternatively, the code can also be compiled for the Cu using the open-source project IceStorm. Finally, the configuration is created and uploaded to the FPGA using AlchitryLabs.

### Computer control

MicroFPGA communicates with the FPGA boards using serial communication, following the register interface protocol developed by Alchitry. The communication interface allows setting and reading various parameters related to the input and output signals. In order to integrate the boards into our microscopes, we wrote a Micro-Manager device adapter that is included in recent Micro-Manager distributions. MicroFPGA properties can be mapped within Micro-Manager to a graphical user interface plugin (20), enabling intuitive control of the different signals and devices downstream from the FPGA. Alternatively, it can be controlled via the provided Java, Python and LabVIEW libraries (18).

### Complementary electronics

We packaged our FPGA within a custom electronics box, together with complementary electronic boards (see Fig. S1 a). Alchitry’s FPGA boards use 3.3 V logic level pins, therefore care should be taken not to input higher voltages. As many hardware devices output or take in 5 V signals, we provide the blueprints for a custom multichannel bi-directional voltage divider (see Fig. S1 b), the signal conversion board (SCB). It can be used to scale up the board outputs to 5 V or scale down digital inputs to 3.3 V in 5 independent channels. Selecting the direction and voltage conversion is performed by bridging the corresponding soldering paths (see Fig. S1 c). It can also be configured to accommodate conversion to an additional custom voltage. This third voltage level is defined by a set of resistors and needs to be chosen prior to soldering. As some devices are controlled with analog signals rather than digital ones, the SCB also features the possibility to low-pass a PWM signal from MicroFPGA, converting it to an analog signal using a Sallen-Key circuit with a cut-off at 10 Hz. Two channels of the SCB feature the low-pass filter. The resulting analog output voltage range is defined by the aforementioned voltage conversion.

As the Au+ board can only measure analog signals between 0 and 1 V, we also provide the description of an 8-channel unidirectional voltage divider for both 5 and 10 V analog signals (see Fig. S1 d), named analog conversion board (ACB). The ACB offers the possibility to limit voltages to 1.6 V, preventing damages to the FPGA ADC. The voltage conversion and limitation can be easily set using jumpers (see Fig. S1 e).

The SCB and ACB are optional and are only needed for particular purposes. If the set-up in which MicroFPGA is installed can work with 3.3 V TTL logic and does not require analog output, there is no need to add an SCB. This is for instance the case if the camera uses 3.3 V signals. Similarly, the ACB is only needed if analog inputs are expected to be used by MicroFPGA. Finally, the electronic box allows more stable connections, but is only optional. Note that the electronic box was designed for the Au+ FPGA using all MicroFPGA inputs and outputs.

### Camera and laser synchronization

Most cameras used in microscopy, whether (scientific) complementary metal–oxide–semiconductor ((s)CMOS) or electron multiplying charge-coupled device (EMCCD), can be synchronized with other devices using TTL signals. Synchronization between the camera and the lasers avoids for instance unnecessary excitation and bleaching of the sample in between two consecutive frames. The nature of the synchronization depends on the camera type and its available trigger modes. For instance, the camera often emits a TTL signal corresponding to the time interval where light is collected by the sensor. In such cases the TTL signal, referred to as the *exposure signal* here (see blue signal in Fig. 2a), is high when the camera is exposing and low when it is registering the sensor pixel values. Alternatively, cameras can be triggered by a TTL input. Each new pulse of the TTL input triggers the acquisition of a new frame. We refer to the TTL input in this manuscript as the *fire signal* (see red signal in Fig. 2b). The exposure time is in some cases set directly by users through an acquisition software, or encoded in the duration of each pulse of the *fire signal*.

**Fig. 2.**
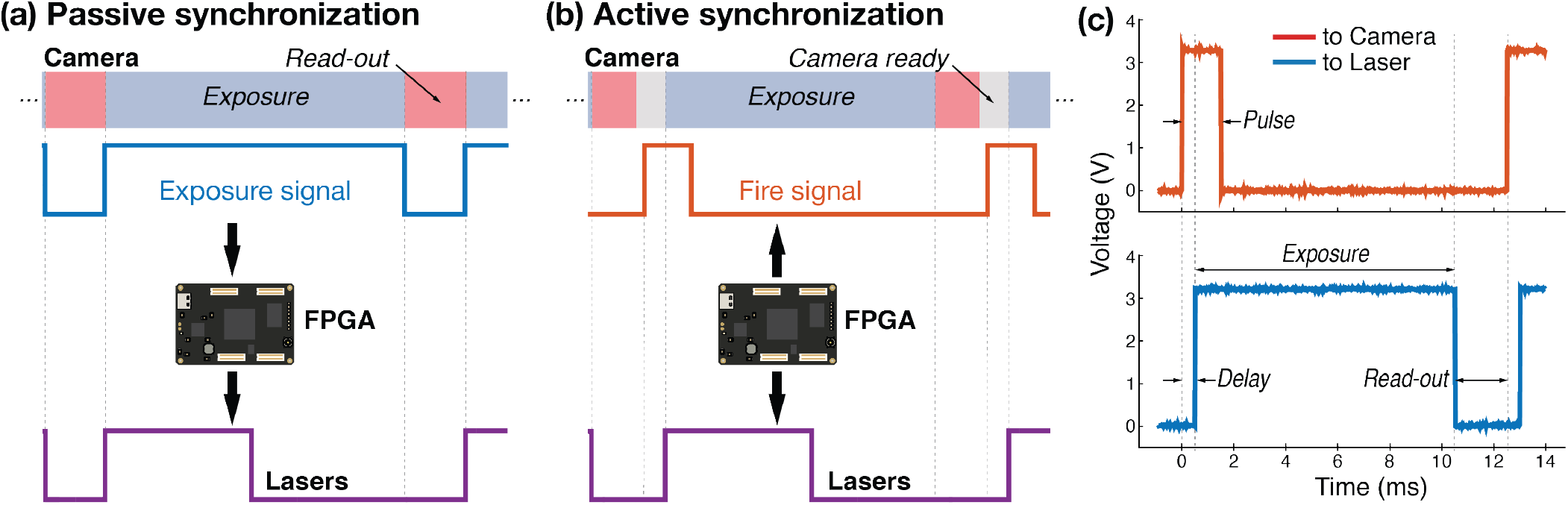
Passive and active synchronization. (a) In passive synchronization, the camera generates an *exposure signal* (blue), which is then processed by the FPGA in order to trigger the lasers (purple). (b) In active synchronization, the FPGA generates a *fire signal* (red) used to trigger the camera. There is usually a delay between the camera receiving a pulse and the start of the next frame acquisition. The FPGA also generates the laser trigger signals (purple), in sync with the *fire signal*. (c) active synchronization requires four parameters: the pulse parameter is the pulse length of the *fire signal* (in red), the delay parameter introduces a delay between fire and an internal *exposure signal* (see red and blue signals), while the exposure parameter is the pulse length of the *exposure signal* (in blue) and the read-out parameter introduces a pause between the end of the exposure and the next fire pulse. We used the following camera parameters: pulse = 1.5 ms, delay = 0.5 ms, exposure = 10 ms and read-out = 2 ms.

#### Camera synchronization

MicroFPGA offers two synchronization modes for the camera: passive and active. In passive mode (see Fig. 2a), the camera generates an *exposure signal* that is processed by the FPGA in order to trigger the lasers. Rather than simply relaying the *exposure signal* to the lasers, as typically done, the processing allows more flexibility in the laser behaviour (see the next section for a description of the laser triggering parameters).

In active mode, the FPGA generates both the input trigger to the camera (*fire signal*) and the laser trigger signals (see Fig. 2b). The *fire signal* shapes will depend on the particular application and camera trigger mode. In order to cover a large range of possibility, the *fire signal* is entirely parameterized by four parameters: pulse, delay, exposure and read-out (see Fig. 2). The pulse parameter corresponds to the pulse length of each pulses in the *fire signal*. The pulse duration can be set to a small value or to the full exposure length, depending on the trigger mode in which the camera is. In active mode the FPGA also generates an internal *exposure signal*. This signal is then processed in the same way as an actual camera’s *exposure signal* in passive mode. Most often, there is a delay between the camera receiving a *fire signal* pulse and the start of the exposure. The delay parameter accounts for such a delay between the *fire* and *exposure signal*. The latter is then pulsing for the duration of the exposure parameter. Finally, the next *fire signal* pulse starts after a read-out period, leaving time for the camera to register the pixel values before the next pulse. This means that the *fire signal* is periodic, with a period equal to the sum of the delay, exposure and read-out parameters. All parameters are set in steps of 1 μs with a maximum value of about 1 s for the pulse and exposure parameters, and 65 ms for the other parameters. MicroFPGA waits for a computer command before generating the signals and similarly stops the triggering upon receiving the corresponding instruction (see the extra parameter in Table 1).

The various parameters should be chosen in agreement with the specification and settings of the camera. For instance, for a camera triggered at every pulses and an exposure length set by the acquisition software, the pulse parameter can be experimentally set by increasing the pulse length until frames are received by the computer. The exposure parameter should reflect the exposure from the acquisition software. The delay and read-out parameters depend on the camera itself, and can be approximated experimentally by observing the pixel intensity in the acquired frames.

#### Laser triggering

Regardless of the camera synchronization mode, MicroFPGA processes an (internal or external) *exposure signal* and allows complex triggering of multiple independent lasers (by default 8). Five different trigger modes are available: on, off, rising, falling and follow. In on and off modes, the laser remains in the same state regardless of the camera trigger. In rising and falling modes, the laser is pulsed at the rising or falling edge of the camera trigger (see Fig. S2 a), respectively. The pulse length in both rising and falling modes can be set using the duration parameter. Finally, in follow mode, the laser follows the *exposure signal* (Fig. S2 a). When triggering a laser, we observe a mean delay of 58.4 ± 2.7 ns (N=203) between the camera and the laser signal (see Fig. 3a) in passive synchronization mode, and 54.3 ± 0.2 ns (N=232) in active mode (see Fig. S3 a). The pulse duration in rising and falling modes is between 0 and 65535 μs, with 1 μs steps. Fig. 3b (passive) and Fig. S3 b (active) showcase an example of three lasers simultaneously pulsed for 1, 2 and 3 μs. Since the FPGA has a 100 MHz clock and fast rise time, the pulses are extremely accurate, with standard deviations on the order of 0.1 ns (N=205 and 208 for passive and active sync modes, respectively). If the pulse duration is longer than the triggering interval of the camera corresponding to the exposure, the pulse length is automatically shortened to this interval.

**Fig. 3.**
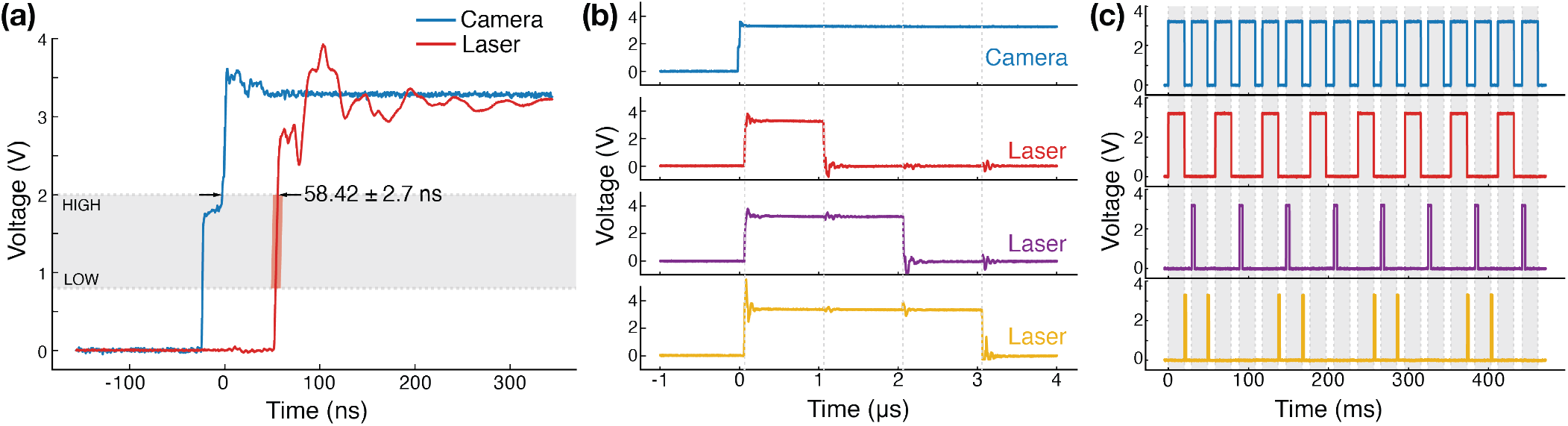
Laser triggering. (a) Measured delay between the camera *exposure signal* (blue) and a laser trigger generated by the FPGA (red, passive synchronization). The laser trigger corresponds to the median delay of 203 signals recorded sequentially. The average delay ± standard deviation is indicated on the plot. LOW (0.8 V) and HIGH (2 V) thresholds are the typical maximum (LOW) and minimum (HIGH) TTL input voltages for the corresponding states. The shading during the transition corresponds to the maximum and minimum observed delays. (b) Multiple lasers can be pulsed in parallel with different pulse lengths, example here with 1 μs (red), 2 μs (purple) and 3 μs (yellow), while the *exposure signal* generated by the camera and processed by the FPGA is shown in blue. (c) Triggering pattern of three lasers using the following parameters: [mode, duration (μs), decimal sequence = binary sequence]. Laser 1 (red) with [follow, not applicable, 43690 = 1010101010101010], laser 2 (purple) with [rising, 4000, 21845 = 0101010101010101] and laser 3 (yellow) with [falling, 2000, 52428 = 1100110011001100]. The camera exposure signal is shown in blue.

All periodic trigger modes (rising, falling and follow) follow a triggering pattern parametrized by a sequence of 16 bits. Each bit represents a camera frame at which the laser is triggered (1) or not (0) (see Fig. S2 b). Since the laser signals share a unique counter, the patterns of all lasers, although independent, are synchronized. Fig. 3c demonstrates the flexibility of MicroFPGA triggering system; the first laser is in follow mode but triggered only every two frames (sequence parameter equal to 43690 or 1010101010101010 in binary), the second laser is triggered on the rising edge (rising, duration equal to 2 ms) with a pattern opposite to the first laser (sequence equal to 21845 or 0101010101010101 in binary), and finally, the third laser is pulsed on the falling edge (falling, duration equal to 2 ms) with the following pattern: 1100110011001100 (sequence equal to 52428). A similar triggering experiment in active synchronization can be found in Fig. S3 c.

Such a flexible laser triggering increases the experimental possibilities on the microscope. In particular, the duration parameter allows fine control of the total laser power per frame (see Fig. S4). For instance, in single-molecule localization microscopy (SMLM) (21–23) an activation laser (such as a UV laser) is commonly used to randomly activate a subset of the fluorescent molecules at each frame. Because the molecules bleach over the duration of the experiments, the pool of molecules that can still be activated decreases over time. In order to optimize imaging speed, this decrease is compensated by increasing the power of the activation laser throughout the experiment. Instead of manually increasing the laser power over hours, the laser power increase can be automated. MicroFPGA allows changing the pulse length of the laser, thus drastically increasing the dynamical range of the activation laser power tuning. We illustrate this in Fig. 4 with a superresolved image obtained with an acquisition of 160000 frames with automated activation. During the experiment, the pulse duration of the activation laser (rising mode) is slowly increased from 0 μs to a user defined maximum of 10 ms (in steps of 1 μs) based on the number of localizations measured at each frame.

**Fig. 4.**
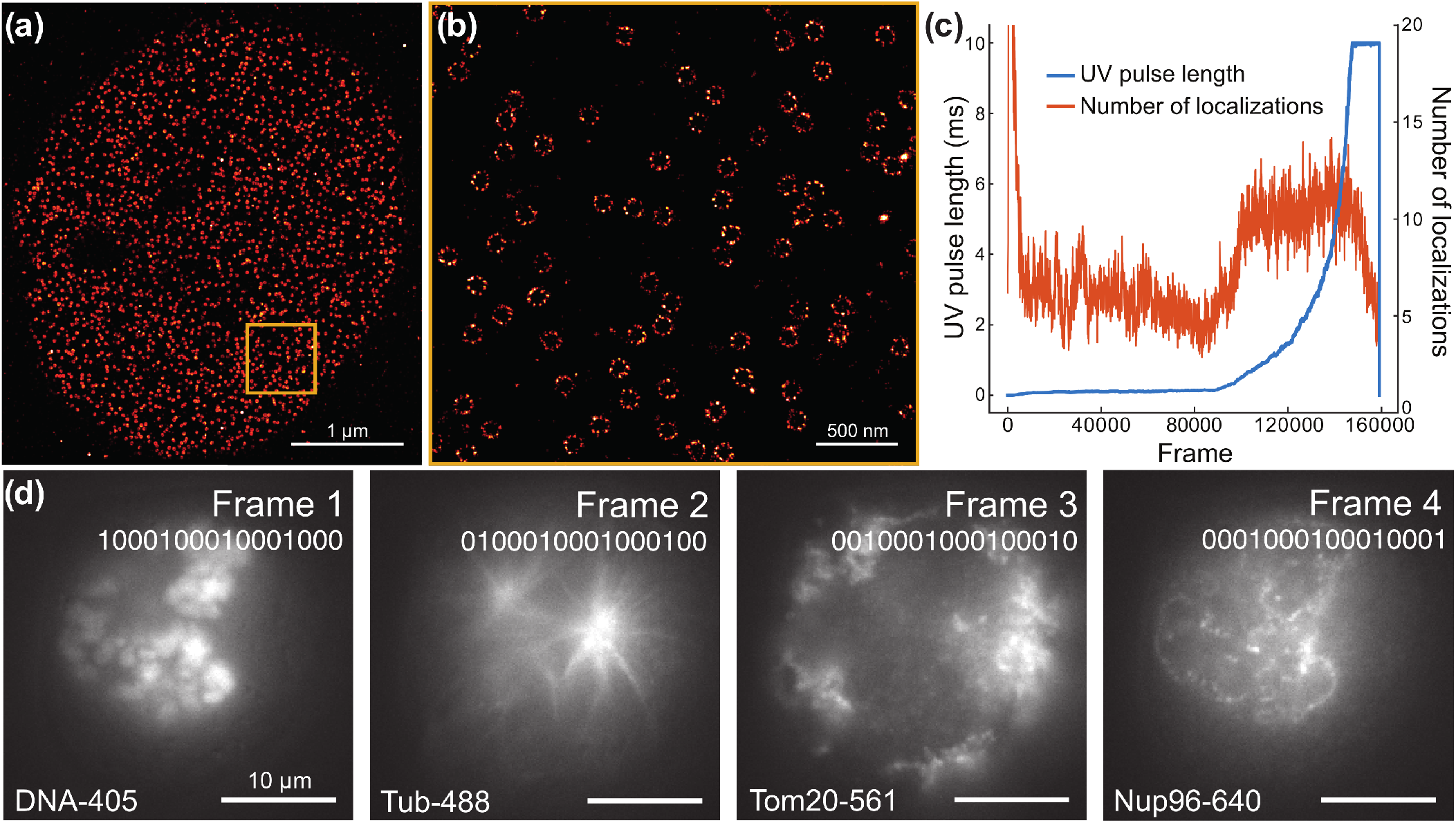
Superresolution imaging. (a) Superresolved image of nuclear pores imaged with single-molecule localization microscopy (160 000 frames) in active synchronization mode. (b) Zoom into the orange region of (a). (c) UV laser pulse length (blue) and number of localizations per frame (red, rolling average of 100 frames) for each frame of the experiment presented in (a). (d) Four consecutive frames of a 200-frame wide-field acquisition are shown side by side. The sample contains four fluorescently labeled structures: DNA (DAPI), Tubulin (immunolabeling, AF488), Tom20 (immunolabeling, CF568) and Nup96 (Halo-JF646), to which corresponds a different laser line each. The laser sequence parameter is shown under the frame number.

The mode and sequence parameters also enable new experiments to be carried out. An obvious application is multi-color imaging by alternating the laser lines during the experiment. For instance, alternating laser wavelengths can reduce cross-talks between channels, or separate in time the different colors when imaging all wavelengths through the same filter. Fig. 4d shows four consecutive frames from the same experiment in which four lasers with different wavelengths are alternating (see each frame’s respective sequence parameter). Each laser excites a different fluorescently-labeled structure in the cell: DNA (frame 1), microtubules (frame 2), mitochondria (frame 3) and nuclear pores (frame 4). A longer frame sequence can be found in Fig. S5, visually illustrating the separation in time of the different structures. To illustrate that the sequence is indeed a 16 bit pattern, we also performed an experiment with three alternating laser lines, while the *16^th^* frame is without any laser (see Fig. S6).

### PWM and analog output

#### PWM

Pulse-width modulation is a type of periodic signal that encodes information in the signal duty cycle, that is to say the percentage of time spent in high signal state during a periodic interval. PWM can be used to directly drive certain devices, such as galvanometers or servomotors. In addition, low-pass filtering a PWM signal results in an analog signal, which enables controlling devices such as acousto-optic tunable filters (AOTF) for instance. By default, MicroFPGA has 5 PWM channels with a period of 1.3 ms and a duty cycle (from 0 to 100%) set using 8 bits (0 to 255).

#### Analog output

When a PWM signal is low-pass filtered, the voltage of the resulting signal is directly proportional to the PWM duty cycle. The period of the PWM channels in MicroFPGA is sufficient to ensure a good low-pass filtering with the SCB board (shown in Fig. S1 b), which requires a PWM faster than 100 Hz and optimally around 1 kHz. Fig. 5a shows the analog output resulting from one of the PWM channel, low-pass filtered and converted to 0-5 V range by the SCB. The signal is shown for different values of the duty cycle (0%, 25%, 50%, 75% and 100%). While the noise introduced by the low-pass filter depends on the duty cycle value, it remains fairly small with standard deviations ranging from 0.005 mV (0% duty cycle) to 0.034 mV (75% duty cycle). For all duty cycle values, the standard deviations are less than 1% of the output voltage. Moreover, the PWM to analog signal conversion is highly linear with only a small offset when fitting a linear curve (*v* = 1.01 * *v_expected_* + 0.05, *R* = 0.99), as shown in Fig. S7.

**Fig. 5.**
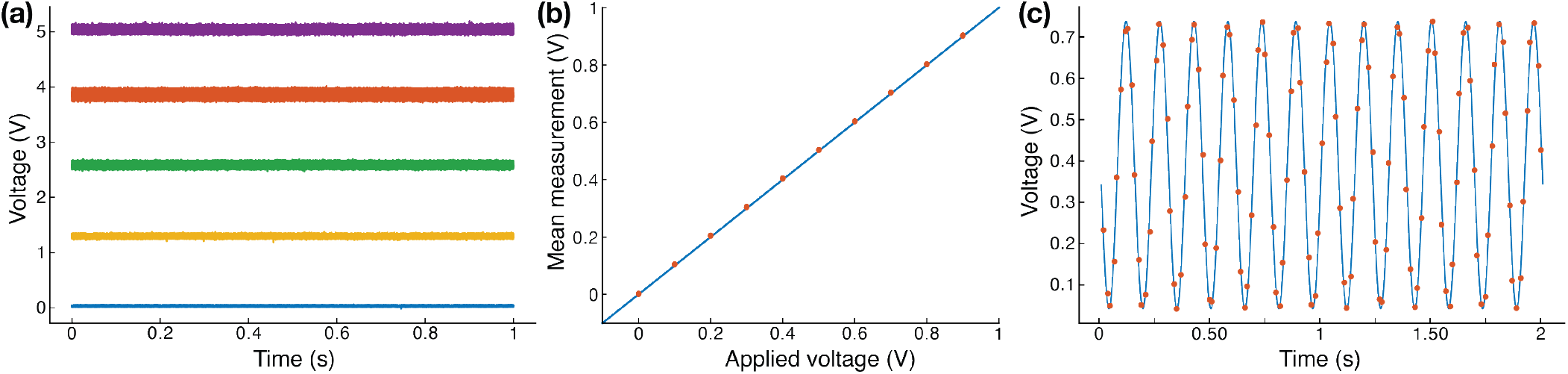
Analog output and input. (a) Analog output voltages measured after passing a PWM signal through the SCB low-pass circuit, corresponding to 0%, 25%, 50%, 75% and 100% of the PWM duty cycle. The average ± standard deviation values are: 0.030 ± 0.005 V, 1.295 ± 0.016, 2.578 ± 0.027 V, 3.856 ± 0.034 V, and 5.046 ± 0.028 V, respectively. (b) Average measurement (red dots, 100 samples each) of the analog input signal normalised to the saturation value (65535) plotted against the applied voltage. The blue line is the expected linear read-out for a perfect analog to digital converter. The standard deviations are comprised between 0.36 and 0.61 mV. (c) Measurements of an analog input channel performed every 16 ms (red dots) and normalised to the maximum value (65535) as compared to the theoretical input signal (blue line), a sine wave of frequency 6.5 Hz.

#### Servomotors control

Servomotors are actuators offering a cost-efficient way to move elements on a microscope (24). For instance, rotary servomotors can rotate a filter wheel or insert a beam block in the optical path, while linear servomotors can serve as stages. Servomotors are controlled by a particular PWM signal whose duty cycle is limited between a minimum and a maximum values that are different from 0 and 100%. These values correspond to the minimum and maximum servomotor positions. In MicroFPGA, the servomotor signal corresponds to the standard signal, which has a period of 20 ms with pulses between 1 and 2 ms. By default, MicroFPGA offers 7 servomotor channels with 16 bits precision for the position (0 to 65535). In order to minimize vibrations resulting from servomotor operation on the optical table, the servomotor signals are switched off 10 s after receiving a new position, allowing them time to move and settle.

### TTL on/off switching

Some hardware devices have the possibility to be switched between two positions (e.g. flip mirrors) or on/off (e.g. lasers) based on an external TTL trigger. We included in MicroFPGA 5 TTL signals that can be toggled between HIGH and LOW states.

### Analog input

Finally, the Au+ FPGA includes a 12-bit analog to digital converter (Zynq-7000 SoC XADC, Xilinx), allowing reading analog signals between 0 and 1 V on multiple channels. Eight such channels are available within MicroFPGA. Fig. 5b demonstrates the high linearity of analog voltage measurements, with standard deviations ranging from 3.6 × 10^-4^ V to 6.1 × 10^-4^ V. Here, the analog input read-out speed is limited by the serial port communication and the transfer between the FPGA and the board microcontroller, the latter being in charge of communication with the computer. To illustrate the board capacity to measure signals in time, we measured a 6.5 Hz sine signal over time (see Fig. 5c). The FPGA board returned a new value on average every 16 ms and the experimental data agreed with the theoretical sine wave (RMSE = 0.03 V).

## Discussion

In this work, we presented MicroFPGA, a FPGA platform that delivers a variety of common signals used to control microscope elements. It is based on affordable FPGAs from Alchitry (Au+ and Cu), and consists of the source-code for the FPGA configuration, as well as complementary electronics to scale signals up or down, and libraries to communicate with the platform from Micro-Manager, Java, Python and LabVIEW.

FPGAs have a clear advantage over microcontrollers in their capacity of processing signals in a parallel fashion and at higher speed. Additionally, increasing the number of tasks carried out by an FPGA does not require timing optimization as long as the tasks are independent and the FPGA has enough free logic cells. The same situation is not true for microcontrollers, where any new task requires careful optimization in order to not cause delays to other tasks. Typically, triggering several lasers in parallel at microsecond scale is a difficult task to achieve with an Arduino, while it poses no hurdle to an FPGA. Nonetheless, for simple tasks or tasks that are not limited by precise timing, Arduino microcontrollers remain the electronic of choice. As an alternative to home-made electronics, ready-to-use commercial solutions exists, such as the Triggerscope (Advanced Research Consulting).

The main difficulty working with FPGAs is the steep learning curve of HDL-like languages. Since MicroFPGA is based on the FPGA boards from Alchitry, it was written in Lucid, a friendly HDL language developed by the manufacturer. In addition, Alchitry provides on their website a set of tutorials that is continuously expanding, lowering the entry barrier to program their FPGAs. Thus, users wanting to modify MicroFPGA can draw inspiration from these tutorials and modify the source-code to fit their applications. To this end, we also uploaded to the MicroFPGA repositories brief tutorials covering a range of useful modifications of the code, such as changing the number of signals available in MicroFPGA, remapping the output pins, creating signals to control more exotic servomotors or changing the parameters’ dynamic range. Advanced users could also repurpose the code for applications requiring a different type of control, such as synchronizing actuators with the camera.

Both the Au+ and Cu FPGA run on 3.3 V within MicroFPGA and care should be taken not to input higher voltages. In the case of the Au+, the analog inputs run on 1.8 V and can be damaged if higher voltages are used. To make MicroFPGA compatible with any input and output voltages, we provide designs for electronic circuits that perform voltage conversion and limit voltages that could damage the FPGA. Both custom electronic boards, SCB and ACB, can be built by electronic workshops or ordered from manufacturing companies. The price of a single board was about 130 (SCB) and 200 euros (ACB). We packaged each FPGA and its corresponding complementary electronics in a single 3D-printed housing, with home-made connection panels. The internal wiring is therefore fixed and labeled connectors on the outside facilitate installation as well as signals troubleshooting. The FPGA box is in this case tailored to our microscopes in terms of number of signals and labels, but should cover wide array of use. We estimate a total price of the electronic box of about 1200 euros, including the SCB and ACB boards, and 16 hours of electronic workshop for the soldering and cabling of the box side-panels. The price can vary depending on the amount of PCB boards ordered, and whether the PCB manufacturer also assembles the components on the boards. Table 2 describes the main components in the parts list.

**Table 2.**
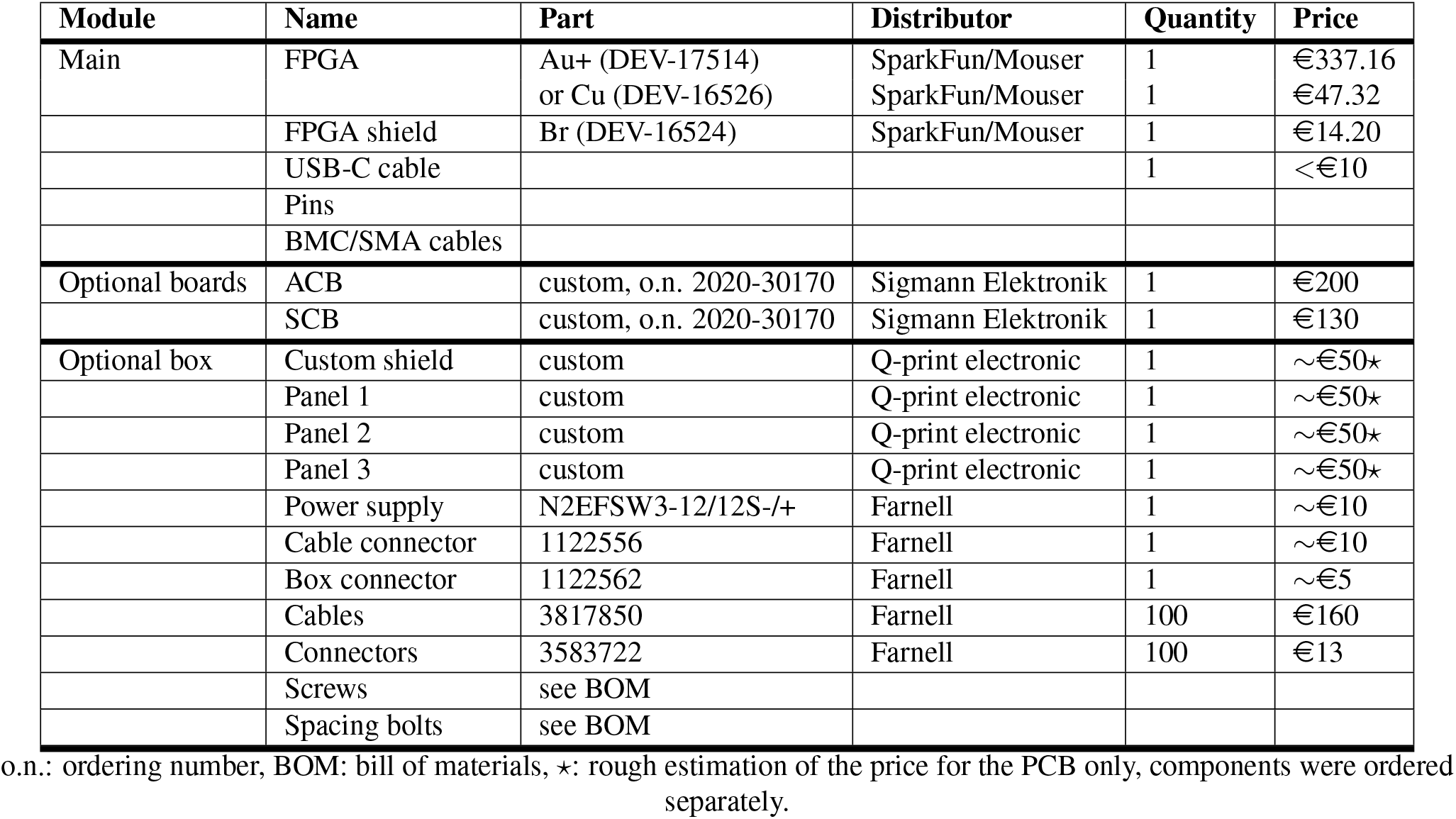
Parts list.

Further improvements to the current design could include shielding the MicroFPGA box to improve signal quality. Likewise, the MicroFPGA code is based on serial communication while the FPGA board is compatible with the open-source USB communication library libusb. The Java and Python libraries, as well as the Micro-Manager device adapters could be modified to use libusb in order to accelerate communication. Finally, new modules could be implemented to make MicroFPGA compatible with galvanometer control or camera ready signals.

MicroFPGA is a key element for the automation of our wide-field microscopes and was tested with many different devices (see Table 3). It is easy to set-up, requiring only access to a soldering iron or an electronic workshop to make the cables necessary to connect devices to the FPGA board. Building the full electronic box requires ordering additional boards with specialised companies or access to advanced electronics facilities. Having a wide range of microscope elements controlled electronically, as it is the case for our microscopes, allows improving the flexibility of the microscope. In particular, the laser triggering allows us to perform localization microscopy experiments with a large dynamical range thanks to microsecond resolution of the activation laser pulses. Since, the laser pulse length can be computer-controlled, we implemented an automatic activation script in Micro-Manager (20), allowing for automated localization microscopy. The patterned triggering also allows interleaved excitation between multiple colors, reducing background and enabling more complex experimental designs. We also control multiple servomotors on the microscopes with MicroFPGA (see Table 3), which includes linear servomotors moving 3D lenses, filter wheels and Bertrand lenses, as well as flip-mirrors. During imaging, our microscope can be switched to different channels (by choosing filters), imaging modalities (2D vs 3D) or illumination (TIRF (25) vs homogeneous illumination (26)) without requiring manual user intervention. MicroFPGA was instrumental in performing high-throughput imaging of biologically-relevant samples (27, 28), illustrating the benefits of microscopy automation.

**Table 3.**
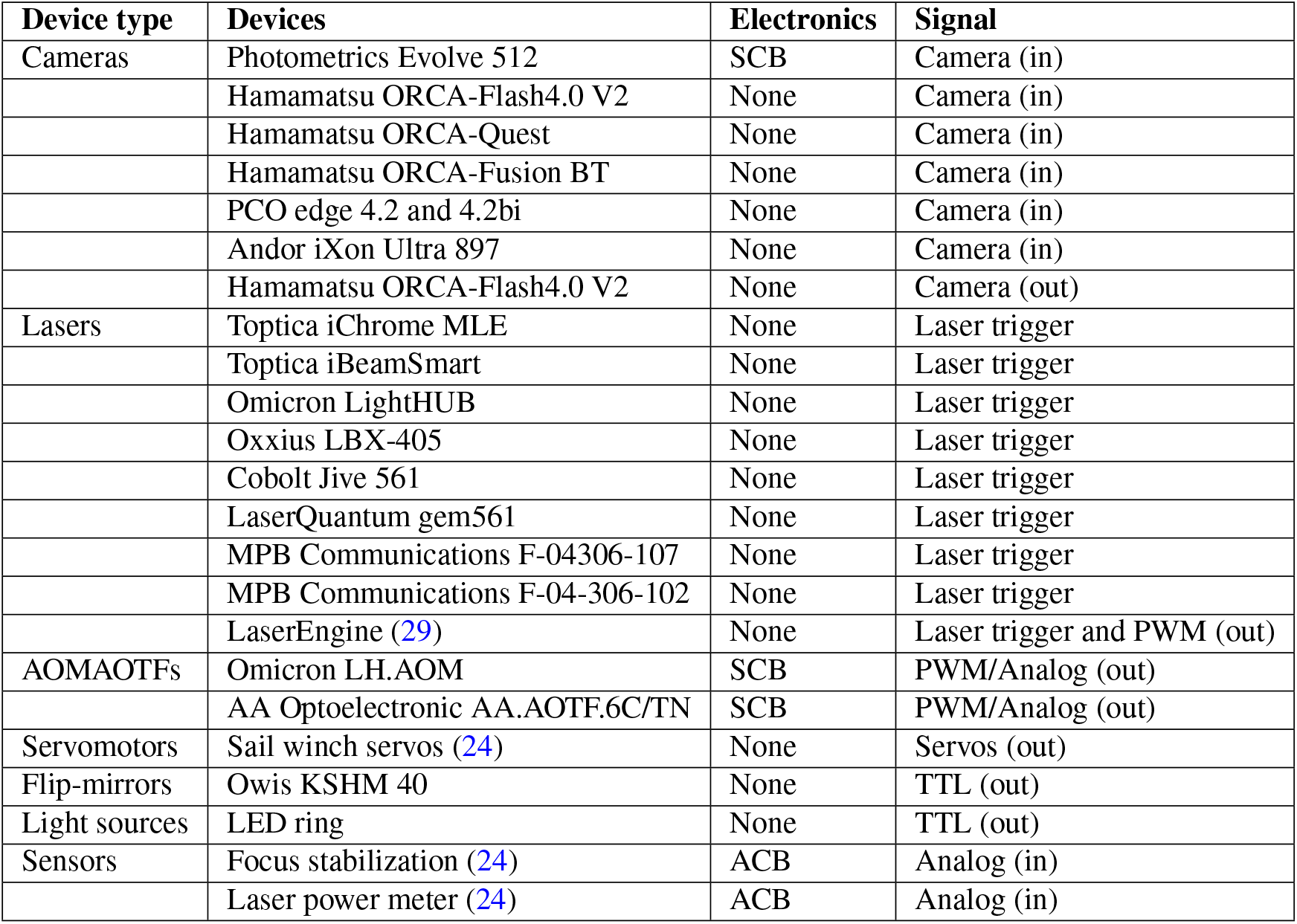
Devices tested with MicroFPGA.

| Device type | Devices | Electronics | Signal |
| --- | --- | --- | --- |
| Cameras | Photometrics Evolve 512 | SCB | Camera (in) |
|  | Hamamatsu ORCA-Flash4.0 V2 | None | Camera (in) |
|  | Hamamatsu ORCA-Quest | None | Camera (in) |
|  | Hamamatsu ORCA-Fusion BT | None | Camera (in) |
|  | PCO edge 4.2 and 4.2bi | None | Camera (in) |
|  | Andor iXon Ultra 897 | None | Camera (in) |
|  | Hamamatsu ORCA-Flash4.0 V2 | None | Camera (out) |
| Lasers | Toptica iChrome MLE | None | Laser trigger |
|  | Toptica iBeamSmart | None | Laser trigger |
|  | Omicron LightHUB | None | Laser trigger |
|  | Oxxius LBX-405 | None | Laser trigger |
|  | Cobolt Jive 561 | None | Laser trigger |
|  | LaserQuantum gem561 | None | Laser trigger |
|  | MPB Communications F-04306-107 | None | Laser trigger |
|  | MPB Communications F-04-306-102 | None | Laser trigger |
|  | LaserEngine (29) | None | Laser trigger and PWM (out) |
| AOMAOTFs | Omicron LH.AOM | SCB | PWM/Analog (out) |
|  | AA Optoelectronic AA.AOTF.6C/TN | SCB | PWM/Analog (out) |
| Servomotors | Sail winch servos (24) | None | Servos (out) |
| Flip-mirrors | Owis KSHM 40 | None | TTL (out) |
| Light sources | LED ring | None | TTL (out) |
| Sensors | Focus stabilization (24) | ACB | Analog (in) |
|  | Laser power meter (24) | ACB | Analog (in) |

The use of MicroFPGA is not limited to custom microscopes, as many commercial microscopes give access to the input and output ports of the cameras and lasers. The FPGA box can therefore be inserted in between to synchronize them. Likewise, some microscopes allow access to parts of the optical path for the purpose of introducing elements such as new filter-wheels, lenses or servomotors. MicroFPGA can be used as long as these elements can be electronically controlled and their inputs are accessible.

In conclusion, we developed an open-source and versatile electronic platform based on an affordable FPGA. The configuration code, Micro-Manager, Java, Python and LabVIEW communication libraries, as well as the blueprints for the complementary electronic circuits, are freely available on Github (18), allowing easy integration of MicroFPGA into existing microscopes.

## ACKNOWLEDGEMENTS

We would like to thank Justin Rajewski (Alchitry) for helping us kick-start the project, and Ulf Matti for his precious help with 3D printing and soldering. This work was supported by the European Research Council (grant no. ERC CoG-724489 to J.R.), the National Institutes of Health Common Fund 4D Nucleome Program (grant no. U01 EB021223 to J.R.), the Human Frontier Science Program (RGY0065/2017 to J.R.) and the European Molecular Biology Laboratory.

## AUTHOR CONTRIBUTIONS

J.D. and J.R. conceived the project. J.D. implemented the code. J.D. and C.K. conceived the electronics. J.D., C.K., P.H. and T.D. performed the experiments. J.D. and J.R. wrote the manuscript with input from all authors.

## Supplementary Note A: Materials and methods

### Programming

The FPGA configuration code was programmed in Lucid within AlchitryLabs 1.2.7 (Alchitry). The Java and Python libraries were developed in IntelliJ 2021.2.3 and PyCharm 2021.2.2 (JetBrains), the Micro-Manager device adapter in Visual Studio 17.0.0 (Microsoft) and the LabVIEW program in LabVIEW 2018 (National Instruments). The FPGA configurations were compiled in AlchitryLabs with iCEcube2 2017.08 (Lattice Semiconductor, Cu FPGA) and Vivado 2020.2 (Xilinx, Au/Au+), before being uploaded to the FPGA using either AlchitryLabs or AlchitryLoader (Alchitry). All source code are available on Github (18).

### Electronics

All electronics boards (ACB, SCB, electronic box side panels, Au/Au+ shield) were designed in Altium Designer V22.3.1 (Altium Limited). All Altium projects were exported to Gerber format and the integrity of the projects was verified using gerbv 2.6A. Both SCB and ACB boards were first hand soldered for testing purposes, while the final versions were ordered assembled from a manufacturer (Sigmann Elektronik) with ordering numbers 2020-30170. The Au/Au+ shield and box side panels were ordered with a PCB manufacturer (Q-print electronic GmbH) and the components were hand-soldered. We tested the various boards individually before assembling the box and before wiring each connection. The electronic box housing was 3D printed in-house and the transparent plastic lid was made by our mechanical workshop.

### Bench measurement

In order to generate Fig. 2c, we connected two channels of an oscilloscope (HDO4054, Teledyne LeCroy) to the camera out (fire signal) pin and the first laser channel of an Au+, through a Br shield (Alchitry). Using a Python script, we set the FPGA to active synchronization and the camera parameters to [pulse (ms), delay (ms), exposure (ms), readout (ms)] = [1.5, 0.5, 10, 2]. We also set the laser parameters (channel 0) to [mode, duration (μs), sequence] = [follow, N/A, 65535]. Since the exposure signal is internal in active synchronization, it cannot be measured directly. A laser in follow mode and sequence = 65535 is the best indirect approximation. We used the oscilloscope to record the two channels for a few tens of milliseconds. Both oscilloscope channels were configured with 1 MOhm impedance.

### Laser trigger: passive synchronization

We used an electronic box featuring an Au board (Alchitry) integrated in a custom microscope, referred here as Microscope A. This microscope has been previously described (30) and includes in particular an EMCCD camera (iXon Ultra 897, Andor) with 3.3 V trigger logic. Because the electronic box is used to control elements on the microscope, it was initially connected to all components (camera, laser, servomotors, filterwheel, bright-field LED ring, flip-mirrors etc.). We added a T-connector on the electronic box in order to measure the exposure signal generated by the EMCCD camera. Since introducing similar connectors on the laser outputs causes voltages to drop, we disconnected the lasers from the electronics and connected instead the oscilloscope channels. Using Micro-Manager (16), we cropped the camera image to 256×256 pixels, set the exposure time to 20 ms and let the camera run live for the duration of the measurements. We used Python scripts to configure the laser parameters; these scripts are available online (18).

For Fig. 3a and b, the oscilloscope was triggered at 2 V on each rising edge of the camera signal and recorded all channels for a short time window (500 ns and 5 μs, respectively). In Fig. 3a, the laser parameters were [follow, N/A, 65535]. We recorded 203 samples. We then used another script to detect the time point at which each signal crosses the minimum HIGH TTL value (2 V) and subtracted the difference between laser trigger and camera thresholds to obtain the delays.

In panel b, the three laser parameters were [rising, 1, 65535], [rising, 2, 65535] and [rising, 3, 65535]. As previously, the oscilloscope was triggered on the camera exposure signal and recorded 205 snapshots containing each laser channel. We detected the rising and falling edges for each laser in order to compute the average and standard deviation of their pulse length. We plotted in the figure the median pulse for each laser.

In Fig. 3c, the measurement was run for 69 frames (about 1 s) and the oscilloscope recorded all four channels (camera and three lasers). We extracted a single window of 16 frames for plotting. The laser parameters were the following: [follow, N/A, 43690 = 1010101010101010], [rising, 2000, 21845 = 0101010101010101] and [falling, 2000, 52428 = 1100110011001100].

### Laser trigger: active synchronization

In Fig. S3, the measurement set-up for the active synchronization was the same as that of the passive synchronization, to the exception of the camera signal. Since in this case the FPGA triggers the camera, we simply connected the first channel of the oscilloscope to the FPGA fire signal output. For panel a, the camera parameters were [pulse (ms), delay (ms), exposure (ms), read-out (ms)] = [1.5, 0, 9, 1], and [1.5, 0, 4, 1] in panel b. The laser parameters were the same as those stated in the previous section. The measurement in panel c had the following camera and laser parameters: [1.5, 0.5, 10, 2], [follow, N/A, 43690 = 1010101010101010], [rising, 4000, 21845 = 0101010101010101] and [falling, 2000, 52428 = 1100110011001100]. The analysis pipeline were the same as before.

### Beads measurement

TetraSpeck beads (0.75 μL; ThermoFisher) with 100 nm diameter were immobilized on a round coverslip (1.5H, 24 mm diameter; Marienfeld) in 400 μL of 100 mM MgCl2 for 5 minutes. After the solution was exchanged for ddH2O, the sample was mounted on Microscope A.

The microscope was controlled by Micro-Manager through an interface plugin (20). We set the FPGA to passive synchronization and used a 647 nm laser (iChrome MLE, Toptica) with parameters [rising, X, 65535], where X is the duration in μs. We used a Micro-Manager script to increase the pulse duration over time between 5 and 500 μs. For each pulse duration, we recorded 50 images of a field of view cropped around a single bead. The images were first averaged for each pulse duration and then average over a region of interest centered on the bead. The curve was fitted with a linear model using scikit-learn (31).

### SMLM experiment

Coverslips were cleaned in a 1:1 mixture of methanol and hydrochloric acid overnight followed by 3 rounds of washing with ddH2O and irradiation with UV. U2OS Nup96-SNAP cells (32) were cultivated under adherent conditions in Dulbecco’s Modified Eagle Medium (DMEM) supplemented with 10% [v/v] fetal calf serum, non-essential amino acids, 2 mM L-glutamine (GlutaMAX), and ZellShield at 37 °C, 5% CO2, and 100% humidity. Cells were seeded 2 days before the experiment on clean coverslips with a density to reach 50 to 70% confluency on the day of the experiment. All following steps were carried out at room temperature. The samples were pre-fixed in 2.4% [w/v] formaldehyde (FA) in PBS for 30 seconds, permeabilised using 0.5% Triton X-100 in PBS for 3 minutes, and fixed in 2.4% [w/v] FA in PBS for 30 minutes on an orbital shaker. Subsequently, they were quenched (100 mM NH4Cl in PBS) for 5 minutes, and washed twice in PBS for 5 minutes each. The samples were blocked with Image-iT FX for 30 minutes followed by 90 minutes of staining (1 μM SNAP-Surface Alexa Fluor 647 [NEB], 1 mM DTT, 0.5% [w/v] BSA, in PBS). Both steps were carried out face down on parafilm in a humidified atmosphere. After the sample was rinsed three times with PBS and washed with PBS three times for 5 minutes each, the sample was mounted on a custom sample holder in imaging buffer (100 U/mL glucose oxidase, 0.004% [w/v] catalase, 10% [w/v] D-glucose, 35 mM cysteamine, 10 mM NaCl, 50 mM Tris-HCl pH 8).

The experiment was performed on another microscope, Microscope B, described previously (33), but with the camera replaced by a different sCMOS camera (ORCA-Flash4.0 V2, Hamamatsu). Microscope B is also controlled using Micro-Manager and the aforementioned interface. The interface plugin includes both an automated activation script for SMLM and an advanced acquisition module allowing to record superresolution images. The automated activation script subtracts two Gaussian-filtered frames and runs a non-maximum suppression algorithm in order to count the number of localizations live. It then updates the activation laser pulse length based on parameters set by the user, among which an approximate number of localization per frame to aim for.

The camera was set to “Global reset level trigger mode” in order to be triggered by the Au FPGA housed in an electronic box in the microscope. The FPGA was set to active synchronization with the following camera parameters: [28 ms, 0 ms, 28 ms, 2 ms]. The lasers were set to follow mode with sequence equal to 65535 (all frames), except the activation laser whose mode was set to rising and with an initial duration of 0 μs. We recorded 160000 frames and stopped the experiment when the pulse duration reached a user defined maximum value.

The raw images were analysed in SMAP (34), using a GPU-accelerated experimental PSF fitting algorithm (35, 36). The superresolved image was also reconstructed in SMAP. The pulse duration at each frame was extracted from the image metadata, while the number of localizations per frame was exported directly from our software. We plotted in Fig. 4c a 100-frame rolling window average of the number of localization per frame to allow interpreting the effect of the automated activation script.

### Wide-field experiments

U2OS Nup96-Halo cells (32) were cultivated and seeded as described above. 2 hours prior to sample preparation, Halo-JFX646 (37) was added to the medium (f.c. 200 nM). All following steps were carried out on an orbital shaker at room temperature, except for blocking and staining steps which were performed on parafilm in a humidified atmosphere at room temperature. The sample was fixed in 2.4% [w/v] FA for 30 minutes, rinsed with 100 mM NH4Cl in PBS, quenched in 100 mM NH4Cl for 10 minutes, rinsed with 0.25% [v/v] Triton X-100 in PBS, and permeabilized with 0.25% [v/v] Triton X-100 in PBS for 30 minutes. Afterwards, it was washed with PBS for 5 minutes and incubated in blocking buffer (2% [w/v] BSA, 0.05% [v/v] Triton X-100 in PBS) for 30 minutes. Then, the sample was incubated with the primary antibodies diluted 1:500 in blocking buffer for 2 hours: mouse anti-alpha-tubulin (Cat# T6074, Sigma-Aldrich) and rabbit anti-Tom20 (Cat# sc-11415, Santa Cruz Biotechnology). Next, it was washed twice with blocking buffer for 5 minutes each followed by three rounds of washing with PBS (1, 5, and 10 minutes, respectively). Afterwards, the sample was incubated with the secondary antibodies diluted in blocking buffer for 1 hour: goat anti-mouse Alexa Fluor 488 (1:200; Cat# A-11001, ThermoFisher) and anti-rabbit-CF660C (1:300; Cat# SAB4600310, Sigma-Aldrich). Subsequently, it was washed twice with blocking buffer for 5 minutes each followed by three rounds of washing with PBS (1, 5, and 10 minutes, respectively). Next, the sample was post-fixed in 4% [w/v] FA in PBS for 10 minutes followed by three washes with PBS for 5 minutes each. The sample was stored at room temperature overnight. Just before imaging, the sample was stained with DAPI (1:10,000 in PBS) for 5 minutes followed by three washes with PBS for 5 minutes each. Finally, the sample was mounted on a custom sample holder in PBS.

We used Microscope A to record sequences of 200 frames with alternating lasers in passive synchronization. In Fig. 4d and Fig. S5, the lasers parameters were [follow, N/A, 34952 = 1010101010101010] (DNA, excited by a 405 nm laser), [follow, N/A, 17476 = 0100010001000100] (Tubulin, 488 nm), [follow, N/A, 8738 = 0010001000100010] (Tom-20, 561 nm) and [follow, N/ A, 4369 = 0001000100010001] (Nup96, 640 nm). In Fig. S6, we used three laser lines with the following parameters: [follow, N/A, 4680 = 1001001001000] (Tubulin, 488 nm), [follow, N/A, 2340 = 0100100100100] (Tom20, 561 nm) and [follow, N/A, 1170 = 0010010010010] (Nup96, 640 nm).

### Analog output

In order to measure analog outputs, we used the oscilloscope directly on the electronic box of Microscope A. We used Micro-Manager to sequentially set the PWM state (channel 0) to 0, 63, 127, 191 and 255. These values correspond to 0%, ~ 24.7%, ~ 49.8%, ~ 74.9% and 100% of the maximum PWM value (255). Within the electronic box, the PWM channel 0 was connected to the first channel of the SCB. This channel convert the signal to the 0-5 V range and low-pass it to obtain an analog output signal. The output of the channel is directly connected to one of the box side panel, allowing to measure the analog output level by plugging the oscilloscope to the box. For each PWM value, we recorded the analog signal for about 1 s. The values are plotted in Fig. 5a, and the same tracks are averaged to produce the plot in Fig. S7. In the latter, a linear model is fitted using scikit-learn.

### Analog input

Fig. 5b was obtained by using a variable power supply (6085, PeakTech) connected to an analog input channel of an Au FPGA. We manually set the voltage level and used a Micro-Manager script to query 100 measurements and export the results to a file. The values were averaged for each voltage level and fitted with a linear model using scikit-learn.

Finally, for Fig. 5c, we used a function generator (UTG9005C, UNI-T) to generate a sine wave with frequency 6.5 Hz and amplitude ~ 8 V. We input the sine signal to an analog input channel of an Au FPGA, and used a Python script to query 100 measurements with a timestamp corresponding to when the query returned. Fig. 5c is obtained by generating a theoretical model of the sine wave using the known frequency and the minimum and maximum measurement values as approximation for the amplitude. We used scipy (38) to optimize the phase of the sine wave on the experimental data.

## Supplementary Note B: Supplementary figures

**Fig. S1.**
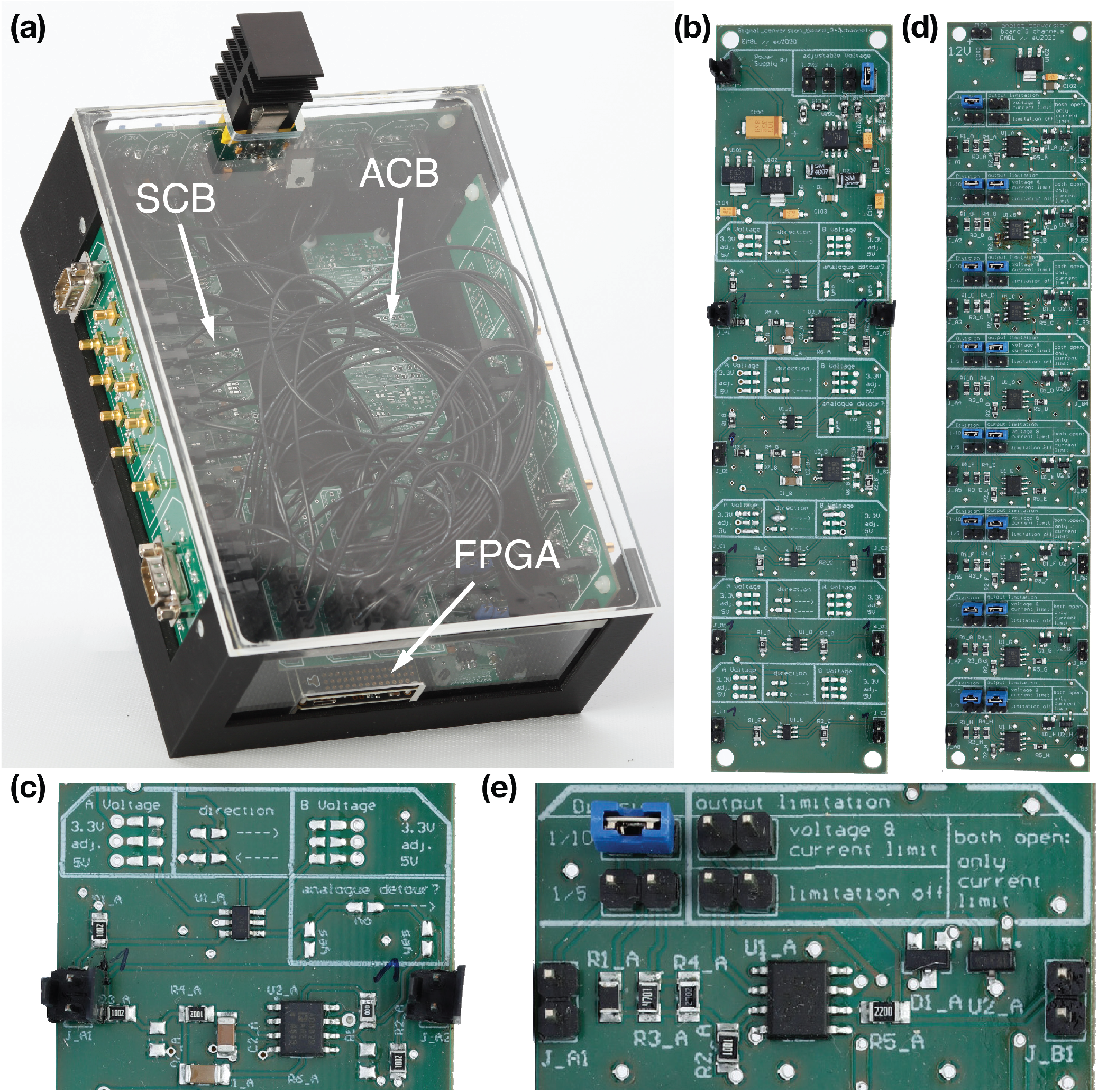
Electronics. (a) Assembled electronics. (b) SCB board. (c) Close-up of the first SCB channel. (d) ACB board. (e) Close-up of the first ACB channel.

**Fig. S2.**
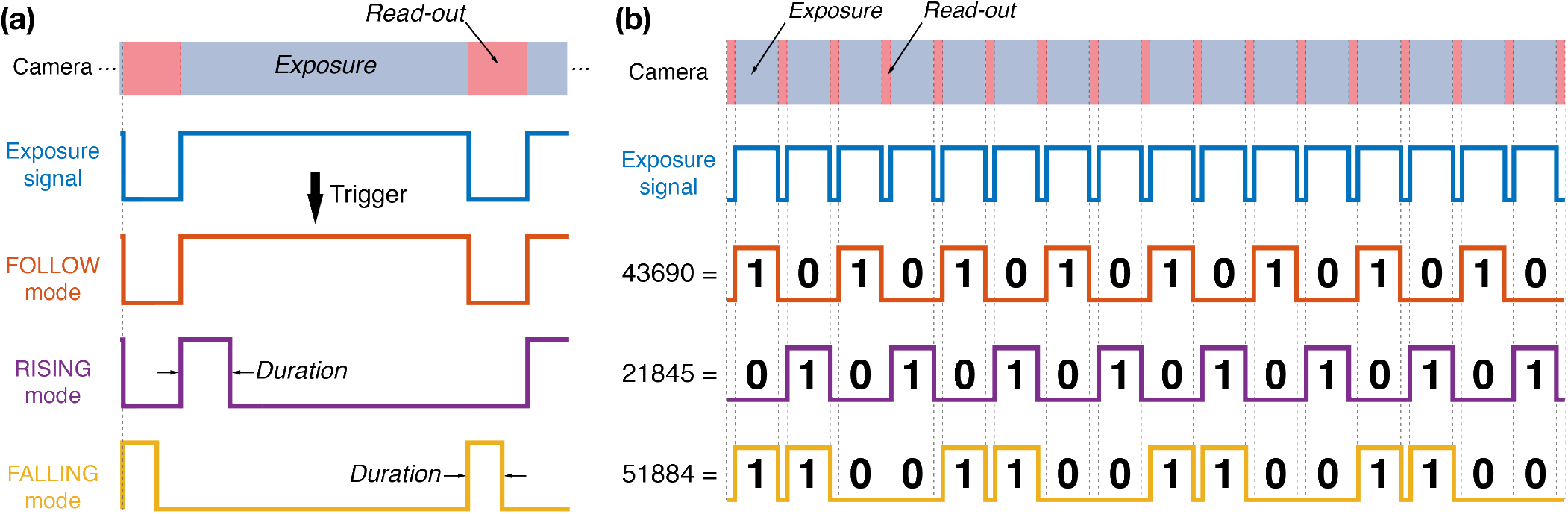
Laser triggering parameters. (a) In passive synchronization, the camera generates an exposure signal (blue) used to trigger the lasers according to different modes: off (not illustrated here), on (idem), follow (red), rising (purple) and falling (yellow). The duration parameter only affect the rising and falling modes. (b) The sequence parameter affects the triggering modes illustrated in (a). It is a sequence of 16 ones and zeroes, each corresponding to the laser being on (1) or off (0) during a frame. The decimal number corresponding to the sequence is shown on the left.

**Fig. S3.**
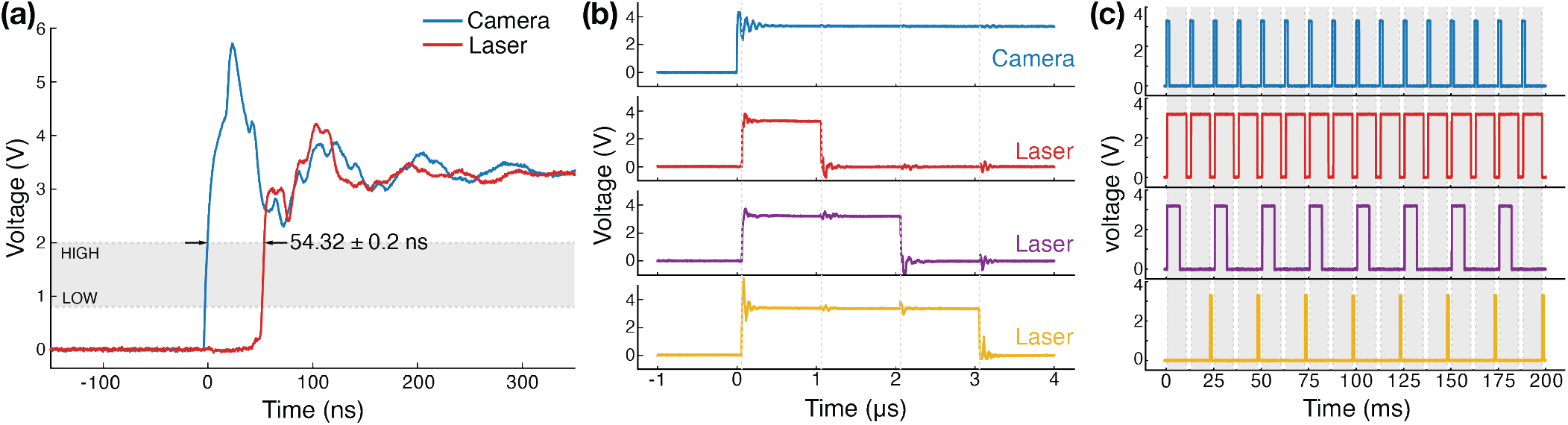
Laser triggering in active synchronization. (a) Measured delay between the *fire signal* (blue) and laser trigger (red), both generated by the FPGA in active synchronization. The laser signal corresponds to the median measured delay of 203 signals recorded sequentially. The average delay ± standard deviation is indicated on the plot. LOW (0.8 V) and HIGH (2 V) thresholds are the typical maximum (LOW) and minimum (HIGH) TTL input voltages for each corresponding state. We did not use any delay (camera parameter). (b) Multiple lasers can be pulsed in parallel with different pulse lengths, here with 1 μs (red), 2 μs (purple) and 3 μs (yellow), while the *fire signal* generated by the FPGA is shown in blue. We did not use any delay. (c) Triggering pattern of three lasers using the following parameters: [mode, duration (μs), decimal sequence = binary sequence]. Laser 1 (red) with [follow, not applicable, 65535 = 1111111111111111], laser 2 (purple) with [rising, 6500, 43690 = 1010101010101010] and laser 3 (yellow) with [falling, 1000, 21845 = 0101010101010101]. The *fire signal* is shown in blue. We used the following camera parameters: pulse = 1.5 ms, delay = 0.5 ms, exposure = 10 ms and read-out = 2 ms.

**Fig. S4.**
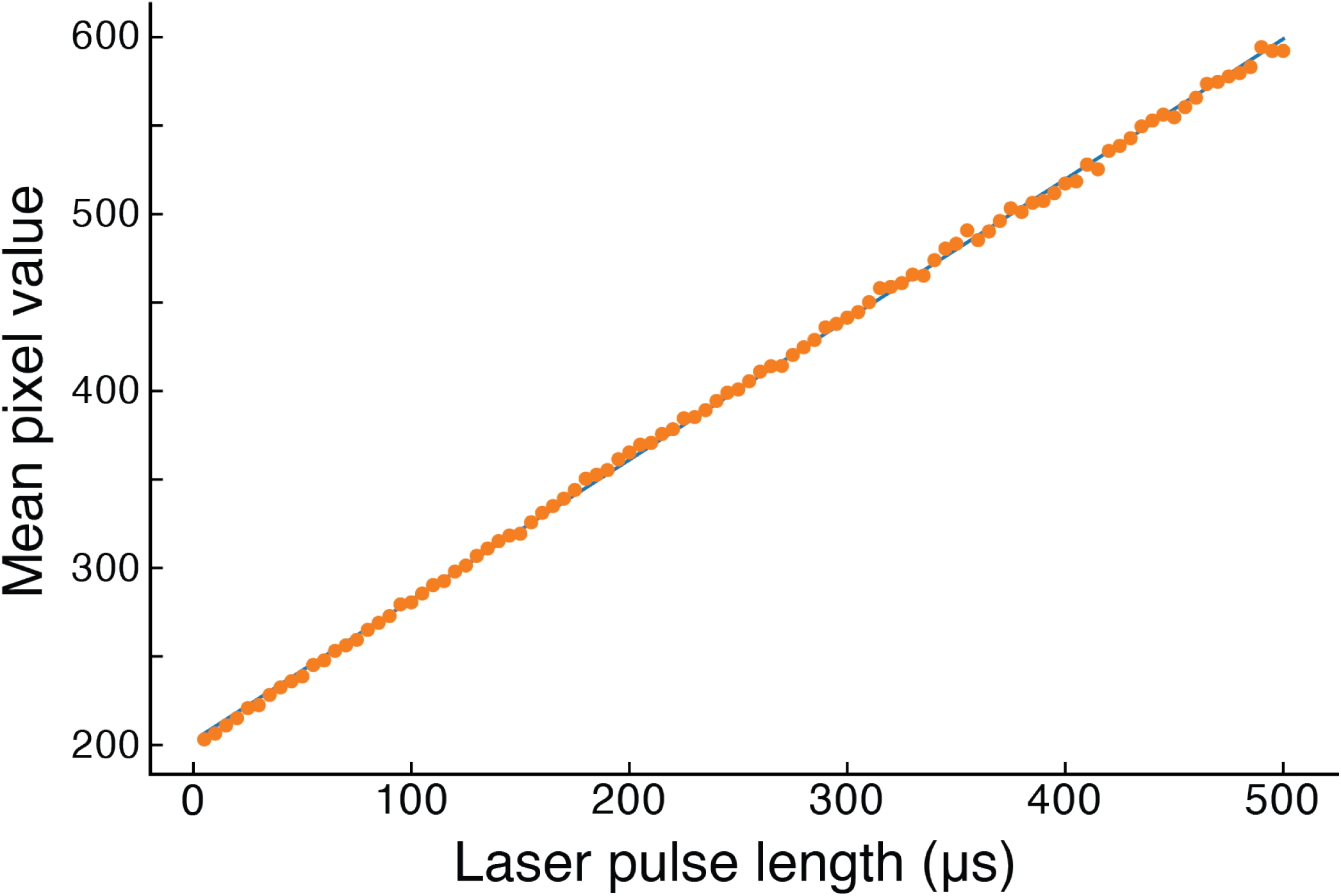
Laser pulse duration and fluorescence. Mean pixel value of a small region surrounding a fluorescent bead plotted against the laser pulse length.

**Fig. S5.**
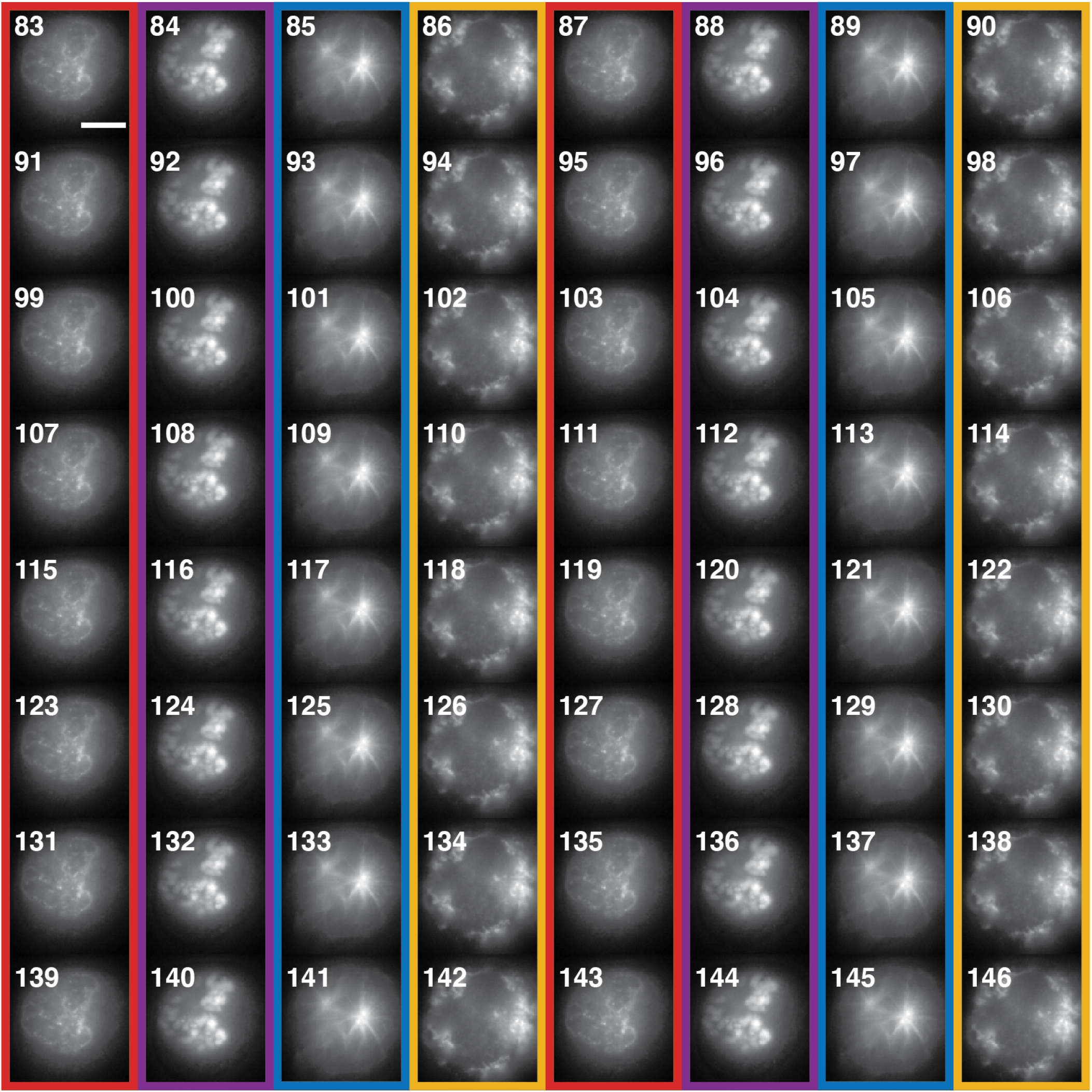
4-color alternating acquisition. Frames 83 to 146 (total of 200) from the acquisition presented in Fig. 3. The columns are colour-coded according to the detected structure: Nup96 (red), DNA (purple), Tubulin (blue) and Tom20 (yellow). The four laser lines are alternating using the sequence parameter of the laser triggering.

**Fig. S6.**
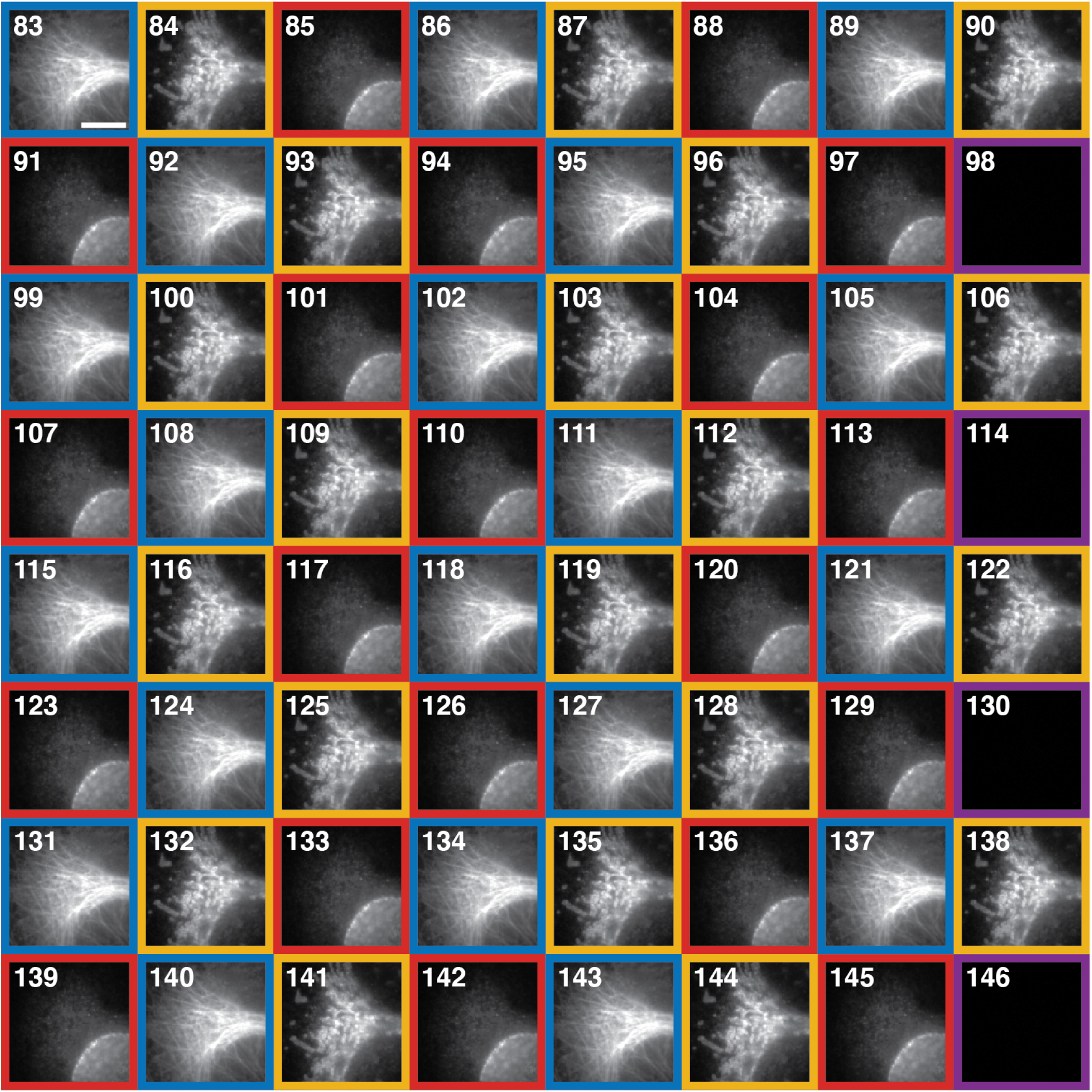
3-color alternating acquisition. Frames 83 to 146 (total of 200) from a 3-color wide-fleld acquisition. The columns are colour-coded according to the detected structure: Nup96 (red), Tubulin (blue),Tom20 (yellow) and none (purple). The three laser lines are alternating. Since the sequence parameter is 16 bits long, the last bit is 0 for all three lasers causing every 16 frames to dark.

**Fig. S7.**
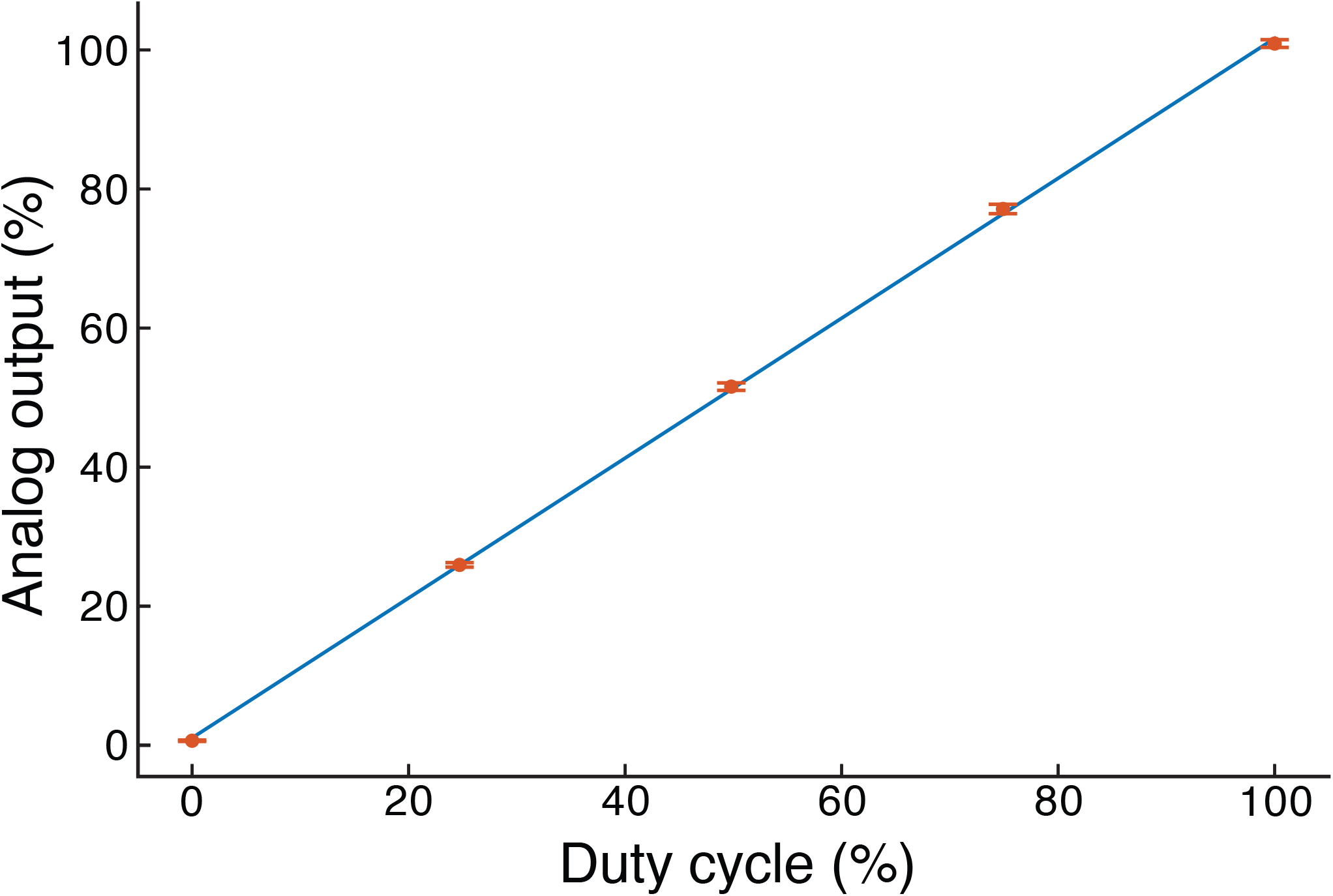
Linearity of the analog output. Mean percentage of the maximum theoretical analog output value (5 V) against the PWM duty cycle percentage (*N* = 10^6^), the error bars correspond to the standard deviation.

## Notes

https://mufpga.github.io/

https://github.com/mufpga

### Competing Interest Statement

The authors have declared no competing interest.

### Summary of Updates

Correct wrong author affiliations and centering of tables.

